# The effect of competency-based education in obstetric emergencies on midwifery students in clinical skill lab, based on Kirkpatrick evaluation model: A randomized controlled trial

**DOI:** 10.1101/695791

**Authors:** Masumah Hakimi, Masoomeh Kheirkhah, Jamileh Abolghasemi, Razia Hakimi, Fatemeh farshad

## Abstract

**Background:** Obstetric emergency is one of the most important causes of maternal and neonatal mortality, and competency-based education is one of the efficient approaches to cover this. Objective structured clinical examination is one of the valid methods in measuring students’ competency and performance. Kirkpatrick evaluation model is a great method to assess a training impact.

**Objectives:** This study was designed to determine the effect of competency-based education on midwifery students based on Kirkpatrick evaluation model.

**Design:** Randomized controlled trial

**Setting:** Nursing and Midwifery School in Islamic Republic of Iran (Iran University of Medical Sciences)

**Participants:** eighty students in third to fifth term of associate and bachelor’s degree in midwifery (intervention group=40, control group=40)

**Methods:** Using stratified random sampling, research team trained learners of intervention group in 4 sessions, 5 hours/day in a month in emergency obstetric cares. Both groups had been receiving the routine schedule of the faculty. Knowledge, skills, and self-confidence were assessed three times, before, immediately and 6 weeks after training by researcher made questionnaire, Objective Structured Clinical Examination (OSCE) and self-reported questionnaire respectively. Data were analyzed with descriptive, inferential statistics.

**Results:** The level of knowledge, skills, and self-confidence increased significantly in the intervention group, in immediate and 6 weeks after intervention (P<0.001). In intervention group, Mean ± S.D of all variables were 5.05±2.074, 143.30±12.146 and 11.65±2.045, which increased to 10.17±1.318, 527.70±19.995 and 18.97±1.980 and remained at the same levels 6 weeks later, 9.37±2.215, 521.80±19.784 and 19.00±2.631; in the control group, this trend was not significant (P=0.380, P=0.455 and P=0.191).

**Conclusion:** Competency-based education can be used in midwifery education and in-service training. We need to use new educational approaches such as competency-based to have a valuable impact on knowledge skills and self-confidence. This may affect health indexes indirectly.

## Introduction

One of the most important causes of maternal and neonatal mortality is the low quality of care provided for mothers and babies which is probably as a result of low levels of skills in health care providers especially in obstetric emergencies. The gap existed between theory and clinical education, can lead to low quality of health services in clinical areas (1). In 2015, almost 303000 women died during pregnancy, delivery and after birth; 830 mothers lose their lives every day because of preventable complications related to birth and 99 percent of them occurred in developing countries (2, 3). In 2017, the maternal mortality rate (MMR) in Iran and Afghanistan were 23.3 and 299.1 per 100000 live births, respectively (4).

One of the most important instructions to constraint maternal and neonatal death is promoting the quality of obstetric care (5). In order to achieve this goal, midwives and gynecologists are being trained. Education using conventional approaches, do not have the required durability; it illustrates the need for using modern teaching methods in the education of midwifery skills (6, 7). The Competency-based educational approach is one of the active and learner-centered methods which increases the ability to perform skills by practicing and repeating (7, 8). Competency in emergency obstetric care is one of the main professional abilities which the International Confederation of Midwives (ICM) emphasizes on it. Shoulder dystocia, bimanual compression of the uterus, manual removal of placenta, postpartum hemorrhage (PPH) management and neonatal resuscitation are some of the common emergencies that achieving competency before confronting them by midwives, can prevent maternal and neonatal mortality and morbidity all over the world (9). Obstetric emergencies teaching in a clinical skill lab can help students to achieve professional competency before clinical environments (10). Learners ‘training by using manikins in a quiet environment before entering the hospital, can decline stress and enhance self-confidence and self-efficacy (11–13). This topic is so important, as more than 65 percent of studies and interventions with the aim of decreasing maternal and neonatal mortality, have been carried out in clinical skill lab or clinical centers (14, 15). WHO and ICM have identified the minimum qualifications, required for midwifery graduates. Competency-based education approach can help to access skill in clinical performance and professional behavior.

To evaluate the effectiveness of this kind of programs, Kirkpatrick evaluation model which assesses the course effect in learners in four levels (Reaction, Learning, Behavior, and impact) can be used. In various studies, different levels of Kirkpatrick evaluation model have been used (16). Since the evaluation of knowledge, skills, and self-confidence in the medical field, especially in midwifery, have a direct relationship with mother and baby lives, therefore it is very important to pay attention to it. This study was conducted to determine the effect of competency-based training in obstetric emergencies on midwifery students in the clinical skill lab based on the Kirkpatrick evaluation model.

## Materials and Methods

### Trial design

This study is a randomized controlled trial with a control group which investigated the effect of competency-based education in obstetric emergencies in midwifery undergraduate students. Using stratified random sampling, learners were divided into two groups by researcher. To control information bias, the evaluation of control group were completed before intervention started. Both groups received regular midwifery emergency obstetric care topics by the faculty members in accordance with the approved syllabus of this country ministry of ministry of health and medical education.

### Participants and setting

All midwifery students in semester 3 to 5, associate and bachelor ‘degrees in nursing and midwifery faculty, informed by posters or directly by the researcher. These students who do not have especial experience in practical skills except internship were eligible to participate. Not participating even in one session was the criterion for exclusion.

### Sample size

To determine the sample size for each objectives, the sample size separately was calculated using the following formula; in which the confidence level (1-α), 95 percent, the test power (1-β), 80 percent, standard deviation (σ) 12.1 and accuracy, 0.5 were considered (17); so the sample size in each group was 40 people.

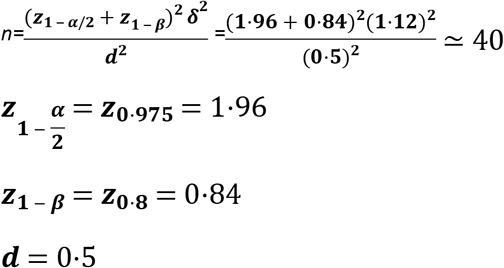

### Intervention

This study was done in accordance to the ethics guideline of Iran University of Medical Sciences (IR.IUMS.REC1397.026) and registered in Iranian Registry of Clinical Trials (IRCT20180609040017N1). All students signed the informed consent. 86 students registered and they were assigned into two groups, intervention (n=43) and control (n=43). First, the objectives and methods of research were explained to learners. All learners completed demographic, knowledge and self-confidence questionnaires. Then, they participated in an OSCE in clinical skill lab based on checklists. The course content was prepared using textbooks and “Emergency Obstetric Care Course Package” of California University. The topics included diagnosis and management of dystocia, postpartum hemorrhage, newborn resuscitation and shock which were taught in the classroom and skill lab center. The validity and reliability of all questionnaire and checklists was verified by 11 midwifery experts and faculty members; the CVI index was generally 0.95 (0.91 - 0.97) and the correlation coefficient using “test-retest” method in the knowledge dimension 0.91, in skill 0.79 and in self-confidence 0.89. Also, the assessors were trained by the researcher and the Kappa coefficient of assessors was calculated at each station which was more than 0.7 in various stations.

The research team conducted the training in 4 sessions which every session duration was 5 hours, once a week, and the assessment was carried out before, immediately and 6 weeks after the end of the intervention. The selected educational approach was competency-based. Firstly, the training began with theoretical topics with different teaching methods such as lectures, watching videos and PowerPoint presentations. The course content in every session was as follows:

Session 1, the reasons for using partograph, filling and interpreting partograph; episiotomy procedure, and its repair, practicing the skill of episiotomy and its repair; Session 2, causes, diagnosis and management of dystopia and its variants, practicing the skill of shoulder dystocia, Session 3, causes, diagnosis and management of newborn resuscitation and shock management in adults; practicing the skill of neonatal resuscitation and shock management, Session 4, causes, symptoms, diagnosis and management of postpartum hemorrhage; practicing the skill of manual removal of placenta and uterine bimanual massage.

In every session, after the theory session, skills were simulated on a model in the clinical skill lab by the researcher. The following checklists were used: episiotomy and its repair, shoulder dystocia, newborn resuscitation, uterine bimanual compression, manual removal of placenta and shock management. Then the learners started group discussion and practicing the skills with supervision. The researcher provided appropriate feedback during practice, and it was continued until competency gained by every learner.

Knowledge assessment was done by a questionnaire consists of 12 multiple choice questions. Every correct answer had one score and the overall score was 12. Self-confidence questionnaire was measured by Likert range of three options from score 1 for “no self-confidence” to 3 for “fully self-confident”. Skills assessed by OSCE in 7 stations. The stations included: Using partograph, episiotomy and its repair, shoulder dystocia, neonatal resuscitation, uterine bimanual compression, manual removal of placenta, and shock management. The assessments were conducted by trained assessors using checklists. Each step was scored by Likert ‘range of 5 options, from score 1 “completely unsatisfactory” to 5 “completely satisfactory”. The total score in each checklist varied based on the number of steps. Cut-off point in knowledge dimension was 80 percent and in skill dimension was 90 percent (18).

### Analysis

Data were analyzed by SPSS version 16. Descriptive statistics such as tables of frequency, numerical indexes and inferential statistics such as Chi-square, Independent t-test, and repeated measures analysis of variance were used to describe the results. The level of significance was considered P<0.05.

## Results

### Characteristics of participants

From 86 participants, (43 students in control and 43 in intervention group), 6 participants excluded from the study; in the control group, 3 students due to illness and unwillingness to continue the study, and in the intervention group, one student due to overlapping the course with her training in hospital and 2 students due to unwillingness to be evaluated had not continued the course and its evaluation. In conclusion, data analysis was carried out on 80 students, 40 participants in each group (table 1: consort diagram).

**Table 1:**
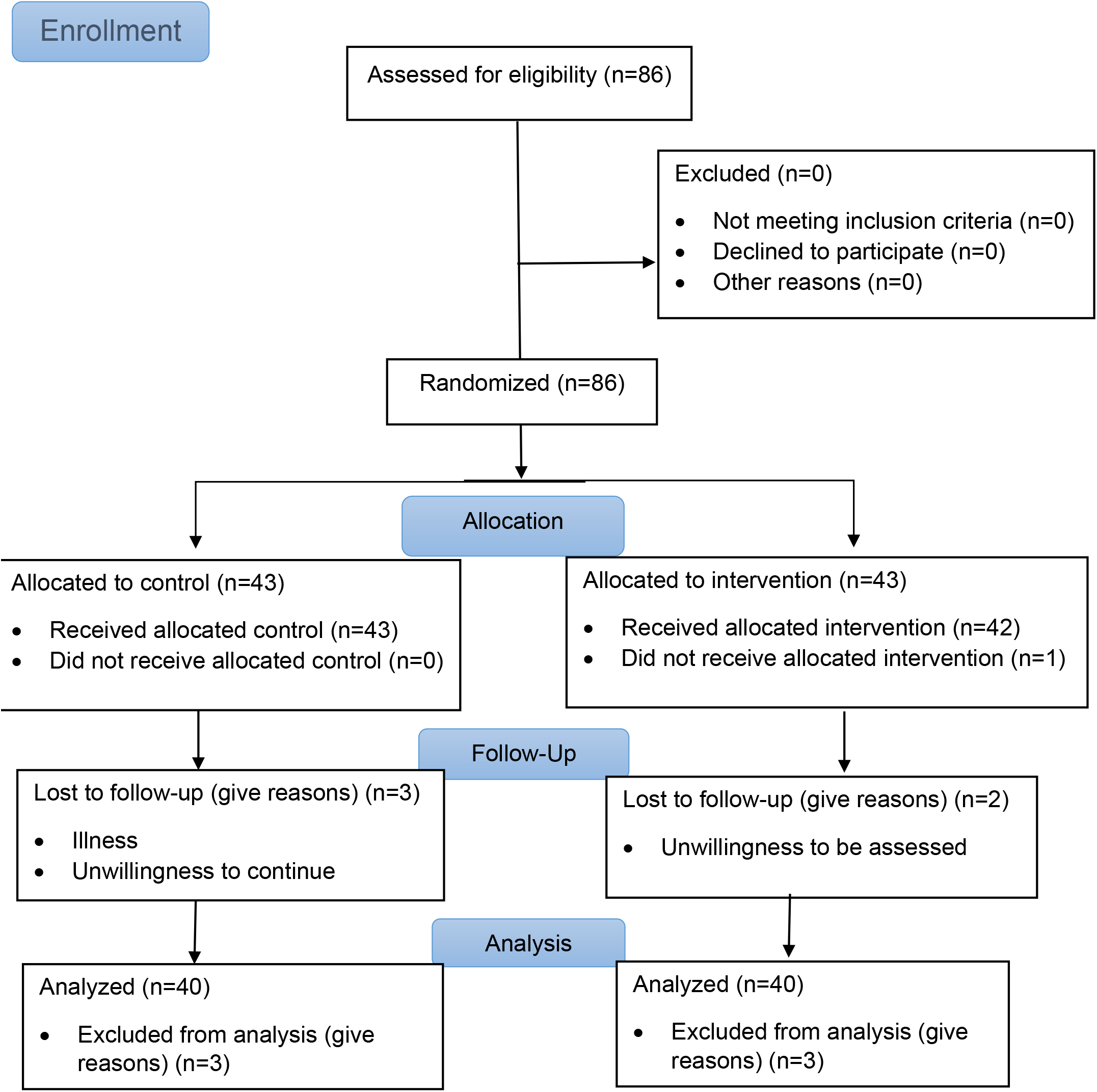
Consort diagram.

Both groups did not have a significant difference in age, residential place, last term average, number of deliveries with coach supervision and independent attendance, experience in dystocia, newborn resuscitation, postpartum hemorrhage and shock management. The majority of students (33.75%) were living in the governmental hostel. Mean and standard deviation of the learner’s age was 21.94 ± 0.475, the last term average was 16.55 ± 0.109. Learners’ demographic features and previous experience in obstetric emergency skills are reported in Table 2.

**Table 2:** Demographic and previous experience in obstetric emergencies workshops in participants.

| variables | Intervention | Control | Statistics |
| --- | --- | --- | --- |
| Age Average ± SD | 21.60 ± 2.38 | 22.27 ± 3.00 | t=1.11; p=0.269 |
| Last term Average ± SD | 16.64 ± 0.97 | 16.47 ± 0.98 | t=0.73; p=0.442 |
| Residential Place |  |  |  |
| Governmental dorm | 16 | 11 | df=2<br>χ²=2.735<br>p=0.434 |
| Non-governmental dorm | 11 | 14 |  |
| Living with parents | 13 | 15 |  |
| Experience in normal delivery |  |  |  |
| Yes | 29 | 32 | df=1<br>χ²=0.620<br>p=0.430 |
| No | 11 | 8 |  |
| Experience in Neonatal Resuscitation |  |  |  |
| Yes | 1 | 2 | p=1.00 |
| No | 39 | 38 |  |
| Experience in PPH Management |  |  |  |
| Yes | 4 | 5 | p=1.00 |
| No | 36 | 35 |  |

### Main outcomes

Knowledge, skills and self-confidence of intervention group increased significantly in immediately and 6 weeks after intervention compared to the control group (Table 3). In comparison between groups, based on repeated measures of variance test, Before the intervention, There were no significant differences in the level of knowledge, skill, and self-confidence among students between groups (respectively P = 0.673, P = 0.153, and P =0.156). The scores of knowledge, skill and self-confidence in all dimensions between groups had a significant difference (P<0.001).

**Table 3:** The effects of intervention on participants.

| Variables | Evaluation Time | Groups (Mean ± SD) | | Confidence Interval | | p-value<sup>1</sup> |
| --- | --- | --- | --- | --- | --- | --- |
|  |  | Intervention | Control | Lower bound | Upper bound |  |
| **Knowledge** | Before | 5.05 ± 2.07 | 4.87 ± 1.58 | 4.29 | 5.63 | 0.673 |
| | Immediately after | 10.17 ± 1.31 | 5.20 ± 1.55 | 4.74 | 10.62 | <0.001 |
| | 6 weeks after | 9.37 ± 2.21 | 5.32 ± 1.50 | 4.72 | 9.97 | <0.001 |
| **p-value<sup>2</sup>** | <0.001 |  |  |  |  |  |
| **Skill** | Before | 118.47 ± 10.14 | 116.22 ± 8.32 | 113.30 | 121.39 | 0.282 |
| | Immediately after | 416.92 ± 19.76 | 116.47 ± 7.07 | 111.80 | 421.59 | <0.001 |
| | 6 weeks after | 415.07 ± 17.28 | 117.90 ± 9.12 | 113.54 | 419.42 | <0.001 |
| **p-value** | <0.001 |  |  |  |  |  |
| **Self-confidence** | Before | 10.52 ± 1.88 | 11.17 ± 1.78 | 9.94 | 11.75 | 0.117 |
| | Immediately after | 16.70 ± 1.75 | 10.75 ± 1.42 | 10.24 | 17.20 | <0.001 |
| | 6 weeks after | 16.75 ± 2.30 | 10.35 ± 2.10 | 9.65 | 17.44 | <0.001 |
| **p-value** | <0.001 |  |  |  |  |  |

<sup>1</sup> Independent t test
<sup>2</sup> Repeated Measures Analysis

In comparison between groups, the score of knowledge, skill and self-confidence in all dimensions, increased significantly in immediately and 6 weeks after intervention (P<0.001). To compare the scores within groups, knowledge, skills, and self-confidence increased significantly from before to immediately, and 6 weeks after intervention, in intervention group (P<0.001); by Bonferroni Post-hoc tests in all dimensions, there was a significant difference between before and immediately and 6 weeks after intervention (P<0.001). There was no significant difference between scores of immediate and 6 weeks after intervention in all mentioned areas, which means that knowledge, skill, and self-confidence were consistent 6 weeks after intervention (respectively P = 0.167, P = 0.464, P = 1.00). Chart 1-3 illustrates the changes and comparisons of knowledge, skill, and self-confidence in intervention and control.

S1-3. Chart 1-3

## Discussion

One of the effective factors in high maternal mortality is the gap of skill performance between theoretical knowledge and clinical activities in “real world”. Based on research team experience, In Iran, despite the appropriate theory teaching which is based on the latest scientific references, midwifery students in a clinical environment do not have the opportunity to achieve experience in obstetric emergency management. University professors train their students in clinical skill lab limitedly and the major part of clinical education is performed in bedside in hospitals. However, they were not allowed to participate in high-risk pregnancy care and management; this can result in dissatisfaction in competency and self –confidence.

Competency-based education is one way to diminish this gap. Also, it is needed to choose an acceptable and efficient method to teach and evaluate skills, first in clinical skill lab, then in the hospital such as simulation and Objective Structured Clinical Examination (OSCE). The ultimate goal of many studies is to upgrade the level of health index in the country in line with Sustainable Development Goals. By using an appropriate design of evaluation like Kirkpatrick Evaluation Model, it is possible to understand the impact of the study on higher levels.

This study was conducted with the aim to determine the effect of Competency-based education of obstetric emergencies on knowledge, skills, and self-confidence of midwifery students with OSCE and using Kirkpatrick evaluation model in its second (knowledge, skills) and third levels (self-confidence). Based on our findings, this study is the first research in the context of educational evaluation with OSCE in obstetric emergencies with a competency-based educational approach in the clinical skill center by using kirckpatric model of evaluation in Iran. All previous studies in these trainings had been done as in-service courses, but our research conducted on students. As half of the students participated in this study were international trainees and they were supposed to return to their country (Afghanistan), it was needed to use a special method which can enhance health indexes such as MMR.

In addition, one of the most precious evaluation method in clinical skills is OSCE that we used in this study. OSCE can generate a beautiful relationship between teaching and assessment. This high valid method can investigate a broad spectrum of clinical skills and individual abilities. Despite the fact that the best place for applying newly taught skills is “real world” and “real patient”, one of the best methods of teaching, practicing and evaluation in some rare urgent activities such as obstetric emergencies, needed to be fulfilled before graduation, is OSCE in clinical skill center (19, 20). The study by Collado-Yurrita at Madrid University approved that utilizing this approach has progressively developed in recent years (21). OSCE investigates the created changes in each individual, but it is clear that it is not enough. The main goal for training is not individual changes, promotion in organization performance is the ultimate goal; one of the foremost methods is Kirkpatrick which assesses the effect of our intervention on different levels of stakeholders such as the impact on the organization, government or country.

Data analysis revealed that knowledge, skill, and self-confidence increased significantly in immediate and 6 weeks assessments after the intervention. The study by Ameh et al. (2018) in which, knowledge and skill retention after emergency obstetric care training with “skills and drills” approach, was investigated, their result was the same as ours; knowledge and skill levels increased significantly immediately and three months after training (P<0.01). The improvement in skill scores was much higher than knowledge and previous job experience more than 13 years caused lower retention and more drop in scores, especially in skill assessment (22). This reduction seems to be due to more skill retention with passing time and increasing experience. This is one of the reasons to start this course during university days, not after graduation or in-service.

This approach leads to competency achievement for all students after the intervention and retention six weeks later in the intervention group and no competency in the control group. In our study cut-off point in knowledge was 80% and in skill 90%. Mirkuzie et al. (2014) studied the reaction and knowledge of health care providers in Ethiopia. 70% of the learners achieved competency in the first assessment after the intervention, and scores remained up until 6 months after training (23). It seems that the course planning after completion of the theory session at university or before starting the internship can have the best impact in results.

The coincidence between teaching and learning education of clinical skills in skill lab by practicing them with a simulation-based approach can improve retention of knowledge and skill as well as self-confidence of students in performing clinical skills. walker et al. study (2015) was conducted with a simulation approach and group practices, an increase and retention of knowledge in a self-report evaluation were observed (24). Their results were in the same line with our study. The results of Walker et al. (2013), showed growth in scores of knowledge and confidence, immediately after intervention; but significant fall within 6 weeks after training (6). Also, the study by Tang et al. (2015) aimed to improve obstetric emergencies ‘knowledge and skills in Malawi also led to an increase in knowledge and skill after intervention (P<0.001), but 6 months after training, both variables decreased significantly (25). It showed that despite the fact that, traditional approaches have this capacity to enhance knowledge and skill levels immediately, but they were not effective knowledge and skill retention. In addition, based on Miller Pyramid, “Know” about a special topic is not enough, and without “Show” and “perform” and practicing skills, we encounter more decay and lower retention. Tang and Walker study results showed that we need to use new educational approaches such as competency-based to have a valuable impact and retention of knowledge and skills and self-confidence.

Our results revealed that using this way of teaching and evaluation can enhance learner’s self-confidence. Increasing self-confidence leads to self-efficacy and as a result, midwives ability in performing skills can grow. Walker et al. study in Mexico resulted in simulation training enhanced self-efficacy followed by better management in emergencies (26, 27). Self-efficacy and self-confidence, especially in emergency case management, are key factors to show competency(8). Studies found that professionals who have more self-confidence and self-efficacy can manage emergency situations better (17, 26).

The results of Glasgow et al. study revealed that a competency-based approach can improve the transparency, efficiency, impact, and quality of education, as well as increasing the accountability of learners and educators (28). KC et al. (2017) also aimed to assess the retention of newborn resuscitation skills after training with Time-series design. In that study, the knowledge and skill scores increased significantly in the immediate evaluation and remained stable until six months after intervention (29). In addition, Urbute et al. (2017) investigated the changes in learners knowledge and self-esteem in obstetric emergencies, with self-assessment and Likert scale (30). Although the results were the same as ours, self-assessment with Likert scale is not an appropriate method to evaluate knowledge and it was one of the items which we pay attention to.

Results of Repeated Measures ANOVA illustrated that knowledge, skills and self-confidence, immediately and 6 weeks after training increased significantly. The Competency-based could enhance the learning outcomes through practice and repetition, especially in medical sciences (21). The results of Ameh et al. (2012) study with Kirkpatrick evaluation method revealed knowledge, and skills increased at the second level and self-confidence at the third level among learners (17). Also, the study by Grace et al in Cameron (2015) was similar to our study. The results showed that knowledge, skills, and self-confidence of learners immediately increased significantly after the intervention. The persistence of knowledge and skills a six-month after the intervention has led to a high level of self-confidence in managing midwifery emergency situations. The theoretical framework of this research was based on Bloom’s theory of education, which divides the educational objectives into cognitive and psychomotor areas; this suggests that using educational approaches which are more likely to sustain the competencies after the intervention, can probably improve the level of self-confidence in managing specific skills. In fact, there is a two-way relationship between these three components, competence, self-confidence, and stress, and all of them are being affected by each other (31). Research by Frank et al. (2009) with 68 interns of obstetrics and gynecology ward, in terms of knowledge and skills, increase immediately after intervention, was in the same line with our study. The results of that study showed that training in a topic like obstetric emergencies using lectures, simulation approach, and instructions of how to use mannequins and pre-prepared scenarios as case study questions, could be effective in increasing knowledge and skills (32). Research by Walker et al. (2014) in Mexico, was in accordance with the present study. The results showed that evidence-based knowledge in the management of obstetric emergencies could be increased with an inter-professional and simulation-based approach, immediately and 3 months after training, followed by self-efficacy. Some evidence suggested that self-efficacy improved as a result of increased self-confidence to diagnose and manage obstetric emergencies (33).

Considering that in-service training has significant costs for the hospital and the government (34), the implementation of this program during university enrollment, after the completion of training and before the start of the internship, could result in a significant reduction in costs. By inserting this course in the midwifery curriculum, and it’s implementation as a part of the midwifery program in the country, as well as conducting in-service training by short-term retraining programs, our peers ‘information can be updated. This intervention was designed to improve the knowledge, skills, and self-confidence of midwifery students in the field of obstetric emergencies. It is recommended that these workshops would be held in codified and approved form in the country.

### Limitations

This study was the results of MSc midwifery thesis and due to lack of time, we could not assess the impact of our study in hospitals. It is recommended that the next study would be conducted in this area.

## Acknowledgments

This article is the result of the master’s thesis of midwifery education at the Faculty of Nursing and Midwifery, Iran University of Medical Sciences with the financial support of the UNFPA Office. The authors wish to thank these organizations, the Italian Embassy, all midwifery students participating in the project, the Center of Clinical Skills in Iran Nursing and Midwifery Faculty, Vice-chancellor for International Affairs and Research of Iran University of Medical Sciences.

